# Origin of division of labor is decoupled from polymorphism in colonial animals

**DOI:** 10.1101/2024.07.05.602267

**Authors:** Sarah Leventhal, Stewart M. Edie, Rebecca Morrison, Carl Simpson

## Abstract

Division of labor, the specialization of sometimes phenotypically divergent cell types or group members, is often associated with ecological success in eukaryotic colonial organisms. Despite its many independent evolutionary origins, how division of labor originates remains unclear. Conventional hypotheses tend towards an “economic” model, so that biological division of labor may reflect a partitioning of pre-existing tasks and morphologies into specialized colony members. Here, we present an alternative model of the origin of division of labor, which can explain the evolution of new functions within a colony. We show that in colonies of the Cretaceous aged (103-96 Ma) fossil bryozoan of the genus *Wilbertopora*, the first cheilostome bryozoan to evolve polymorphism, new member morphologies were not a simple partitioning of pre-existing morphologies, but instead expanded into novel morphospace as they lost functions, specifically feeding. This expansion into new morphologies occurred primarily during two pulses of heightened morphological disparity, suggesting that the evolution of polymorphism corresponded to relaxed constraints on morphology and perhaps to the exploration of novel functions. Using a simple model of physiological connections, we show that regardless of the functionality of these new colony members, all non-feeding members could have been supported by neighboring feeding members. This suggests that the geometric constraints and physiological connectedness could be prerequisites for evolving both polymorphism and division of labor in modular organisms, and that a classic partitioning model of specialization cannot be broadly applied to biological systems.

**One Sentence summary:** In cheilostome bryozoans, polymorphism evolved through the loss of preexisting functions, rather than the gain of new functions, suggesting that polymorphism can evolve through drift rather than division of labor.

## Introduction

The separation of tasks and morphologies among individuals within colonial eukaryotic life (termed division of labor) is widespread today, suggesting that this way of life confers many ecological advantages. Division of labor has evolved repeatedly in both marine and terrestrial organisms, and has been tied to the ecological successes of disparate clades including ants, bees, siphonophores, hydrozoans, and bryozoans (1–4). Specialization of tasks among colony members is thought to increase the energetic efficiency of the entire colony, and thus becomes an emergent feature for selection to act on at evolutionary scales(5, 6). Conventional biological hypotheses for the evolution of division of labor parallel certain principles of economics, where pre-existing tasks once performed by generalists become partitioned among specialists(7, 8). However, this model of partitioning does not explain how the many emergent features of colonial and eusocial division of labor evolved from their ancestral states: bridge-making in ants (9) and the anti-predatory bird’s head avicularia of the bryozoan *Bugula* (10) are both examples of behaviors or functions that are not present in their monomorphic ancestors. Here, we build and test a model of functional and morphological expansion against the classic partitioning model to show that there are multiple, viable pathways for the evolution of division of labor.

There is mixed evidence for the order in which functional and morphological change occur in the evolution of division labor, as the evolutionary sequence has never been directly observed.

Modern eusocial insects are thought to undergo functional change before morphological differentiation (2, 11), while at molecular levels, genetic variation typically precedes functional innovation (12). For example, in gene duplication, one copy maintains ancestral functions while the other can evidently accommodate mutations (and thus eventually accrue new functions) (12). Modules in colonial animals may show similar dynamics, where some members of the colony maintain ancestral functions (13) and others see increased variation while maintaining viability owing to physiological connections with the rest of the colony (direct gut-to-gut connections in bryozoans or communal feeding in ants) (10, 14). To test the relative contributions of functional and morphological change to the evolution of division of labor, we couple direct observations on the functional and morphological variation in an evolutionary time series of a fossil bryozoan lineage with a model of energetic flux. We show that division of labor is not simply a pattern of specialization on pre-existing functional or morphological variation, and that physiological connections in ancestral colonies may be a prerequisite for this evolutionary pathway, as they permit colonies to share surplus energy with non-feeding members.

### Testing evolutionary models for the division of labor in bryozoans

Many cheilostome bryozoans today have evolved division of labor, where their ‘zooids’, or colony members, perform specialized tasks within the colony and express different morphologies, termed polymorphism (3, 14–17). The most common polymorphs include: *autozooids* that feed and produce gametes, *ovicells* that brood embryos (18), and *avicularia* that perform varied tasks from defense to hygiene ([13], Fig. 1). The evolutionary success of cheilostomes through the Cenozoic has been tied to the development of these polymorphs in the Cretaceous genus *Wilbertopora*, 103 Myr ago (19, 20), when avicularia emerged from autozooids, and ovicells formed as composite structures from spine-like zooids (21–23). Thus, fossil colonies of *Wilbertopora* offer critical empirical evidence for these early evolutionary steps towards division of labor in cheilostomes.

**Figure 1.**
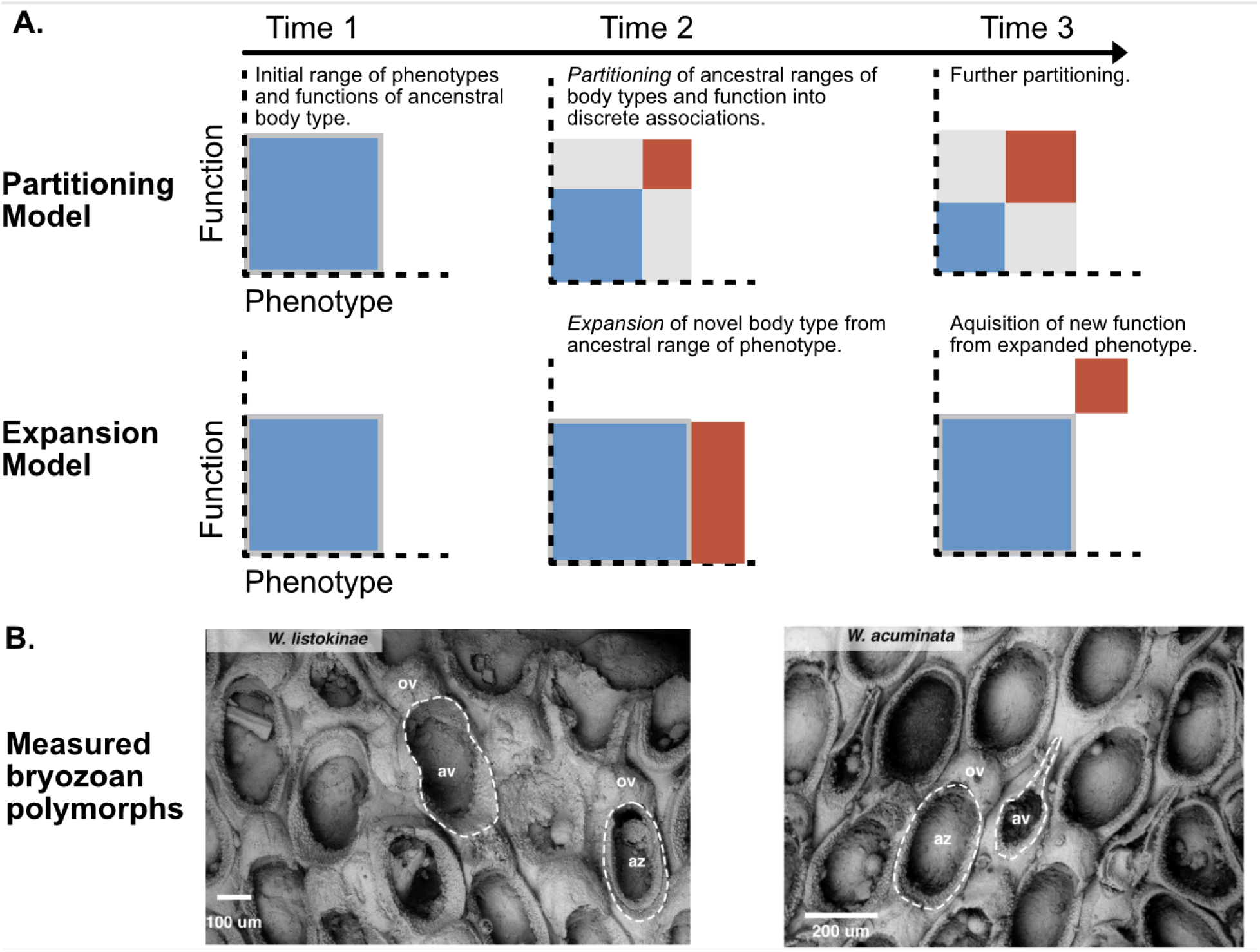
**A.** Two potential patterns of phenotypic and functional evolution in colonial organisms. In the case of *Wilbertopora*, blue squares represent the ancestral body type (autozooids); red squares represent the novel polymorph (avicularia). In the partitioning scenario, avicularia evolve through the partitioning of preexisting functions and variation in *Wilbertopora*. The range of autozooid functions and morphologies decreases as avicularia take on particular functions that used to be performed by autozooids. The total amount of variation in a colony remains similar to that of the ancestral state, but this variation becomes split between autozooids and avicularia. In the expansion scenario, avicularia evolve as an extension of autozooidal variation, expanding the range of variation in a colony. Avicularia then lose autozooid functions and become adapted to completely new functions in the colony. **B.** Polymorphs within two Cretaceous species of *Wilbertopora*. *Wilbertopora listokinae* (USNMPAL 216175), showing an autozooid (az) and avicularium (av) with ovicells (ov). *Wilbertopora acuminata* (USNMPAL 216143).

Polymorphs in extant bryozoans can have discrete and continuous differences in both the types and ranges of their morphologies, which reflect differences in the mode and range of their functions (14, 24). In extant colonies and lineages, autozooids and ovicells have constrained morphologies, corresponding to their functional requirements—avicularia show greater morphological variation, perhaps reflecting the many tasks they can perform (25). Therefore, we measured differences in the morphologies of *Wilbertopora* polymorphs to test whether colonies in this genus more closely followed a model of partitioning or expansion as they evolved. Under the partitioning model, avicularia would have evolved morphologies within the range of autozooids, which would have then decreased in disparity. Under the expansion model, avicularian morphology would have evolved away from the autozooidal range (Fig. 1), which may or may not have also involved changes to the disparities of either polymorph. We can further constrain the effect of shared functions on the evolutionary morphology of in these early colonies by comparing the morphologies of avicularia that lacked the ability to support ovicells to those that had functioning ovicells—these avicularia would have had a functional lophophore and therefore the ability to feed and produce gametes (22) (See discussion in Supplemental Text S7). As avicularia lose these functional ovicells within *Wilbertopora*, we would expect colonies to show increasing morphological differences between avicularia and autozooids.

Beyond the potential morphological consequences of this functional release, the loss of functional ovicells on avicularia potentially presents a different, metabolic challenge to the later colonies of *Wilbertopora*. The inability to feed requires support from the rest of colony, and to test whether an expansion model for the division of labor can accommodate a period of loss of function for members of the colony, we model the energy flux through a set of different colony arrangements to determine if, in theory, there would have been enough energy to support all colony members if only a subset were contributing to energy acquisition. We expect that colonies with a lower fraction of feeding members will necessarily show a decline in their total potential energy acquisition, but the contribution of each feeding member to the energetic output of the colony should increase. We discuss the implications of the new, multi-serial network of zooids on the energy flux for colonies of *Wilbertora* compared to uniserial forms. In combining these models of energy flux with the analyses of morphological and functional change, we present the fossil record of *Wilbertopora* bryozoans as one of the first empirical tests of the different models for the evolution of division of labor.

## Results and discussion

### Evolutionary morphology of avicularia

Consistent with the expectations of the expansion model, zooid polymorphs of *Wilbertopora* evolved increasingly distinct morphologies through time, with avicularia entering new regions of morphospace relative to autozooids both for the genus as a whole and within individual colonies (Fig. 2A-B, Table S3, Table S4; modeled estimates of shape divergence with 95% confidence intervals shown in Fig. S5). Changes in zooid shape, not size, primarily drove this morphological divergence (Table S1, Table S2). Differences in polymorph shapes were mostly derived from changes in avicularian morphology, given that autozooids maintained a similar position and volume of morphospace through the study interval (Fig. 2, Fig. S3). The increased shape divergence between autozooids and avicularia may be underlain by the evolution of new species, as many of the colonies occupying distinct regions of the morphospace are described as distinct species (inferring evolutionary transitions is a reasonable hypothesis, but note that fossil bryozoan species are delimited by the shapes of their polymorphs) (22) (Fig. S6, Fig. S9).

**Figure 2.**
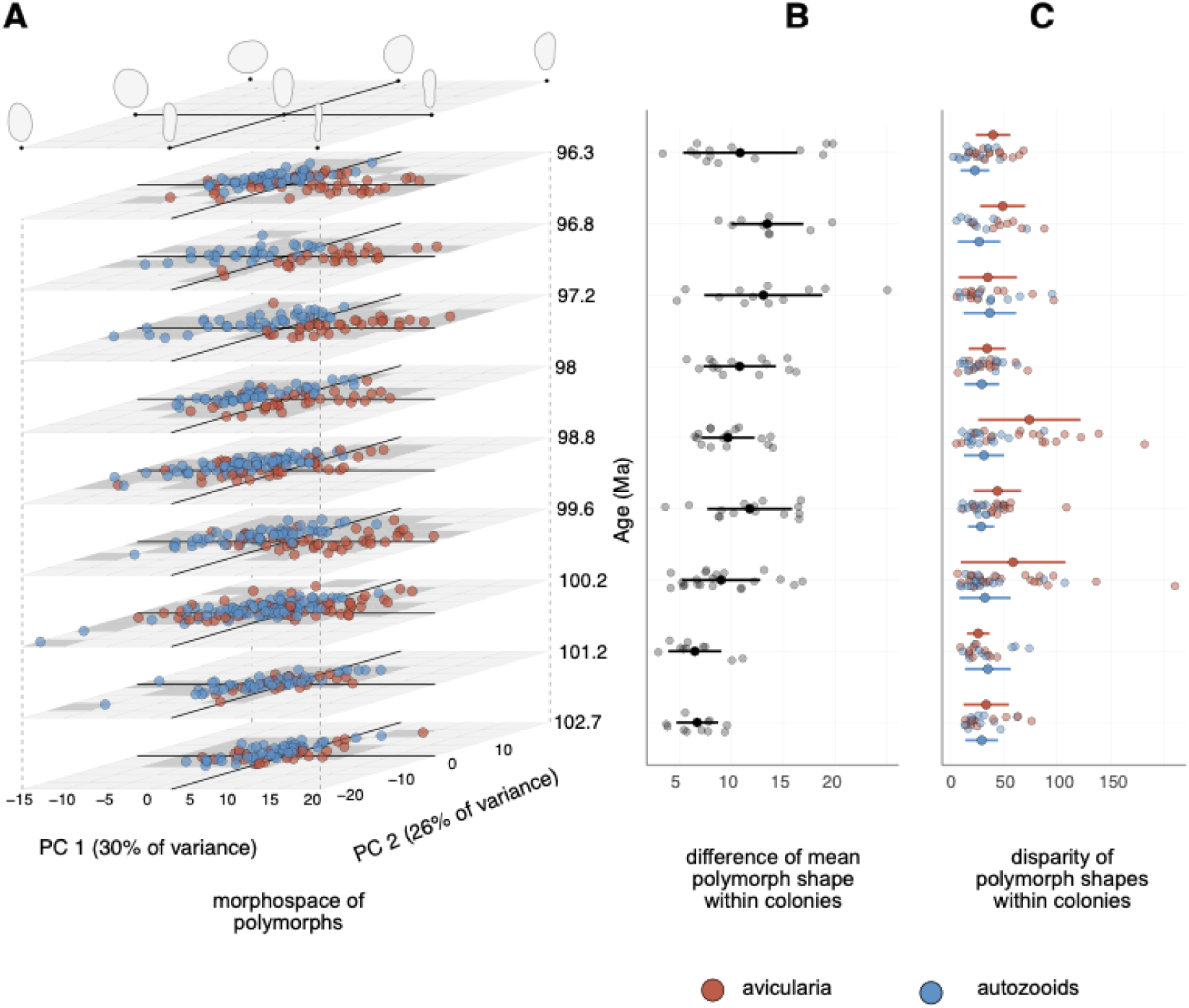
Morphological evolution of polymorph shape (as outlines of a zooid’s orifice and opesia, which together comprise the frontal area of a zooid) in colonies of *Wilbertopora*. **A.** Morphospace of polymorphs defined by first two principal components (PCs). Each point represents a single zooid. **B.** Difference in mean shapes of autozooids and avicularia within colonies (gray points). Black points and bars show mean and standard deviation of values, respectively, within time intervals. **C.** Shape disparity of autozooids and avicularia within colonies (blue and red points); solid points with bars show mean and standard deviations among colonies per time interval. Time slices show the midpoint of stratigraphic formations, but their relative spacing is uniform for visualization purposes.

Regardless of whether these shape divergences resulted from species-level evolutionary events, the net result is an expansion of the earliest expressions of zooid shapes, not a simple partitioning of early variation.

Within the expansion model, it is possible for both the old and new polymorphs to show reduced morphological variation, possibly reflecting specialization on fewer tasks. However, the *Wilbertopora* fossil series shows a general similarity and stability in the morphological variation of zooids within colonies through time, but with two notable exceptions (at 100.2 Ma and 98.8 Mya; Fig. 2A,C, Fig. S7, Fig. S8). In the first half of the time series, two pulses in avicularian disparity correspond to the acquisition of novel elongate forms and the retention of the ovate autozooidal-like forms among colonies (Fig. 2A,C). Neither of these pulses is likely to be an aberration within the dataset, as they do not coincide with known environmental or lithologic changes (i.e. ocean anoxic events (26, 27) and facies changes across formations (28)). The subsequent decline in avicularian disparity to pre-pulse levels corresponds to the decrease in frequency of autozooidal-like forms, and consequently the shift in shape divergence between these two polymorphs (Fig. 3). Thus, *Wilbertopora* evidently fits the expansion model for the evolution of its polymorphs, with avicularia expressing novel morphologies and higher levels of within-colony morphological variation than autozooids, suggesting that changes in the function of avicularia may have also been underway.

**Figure 3.**
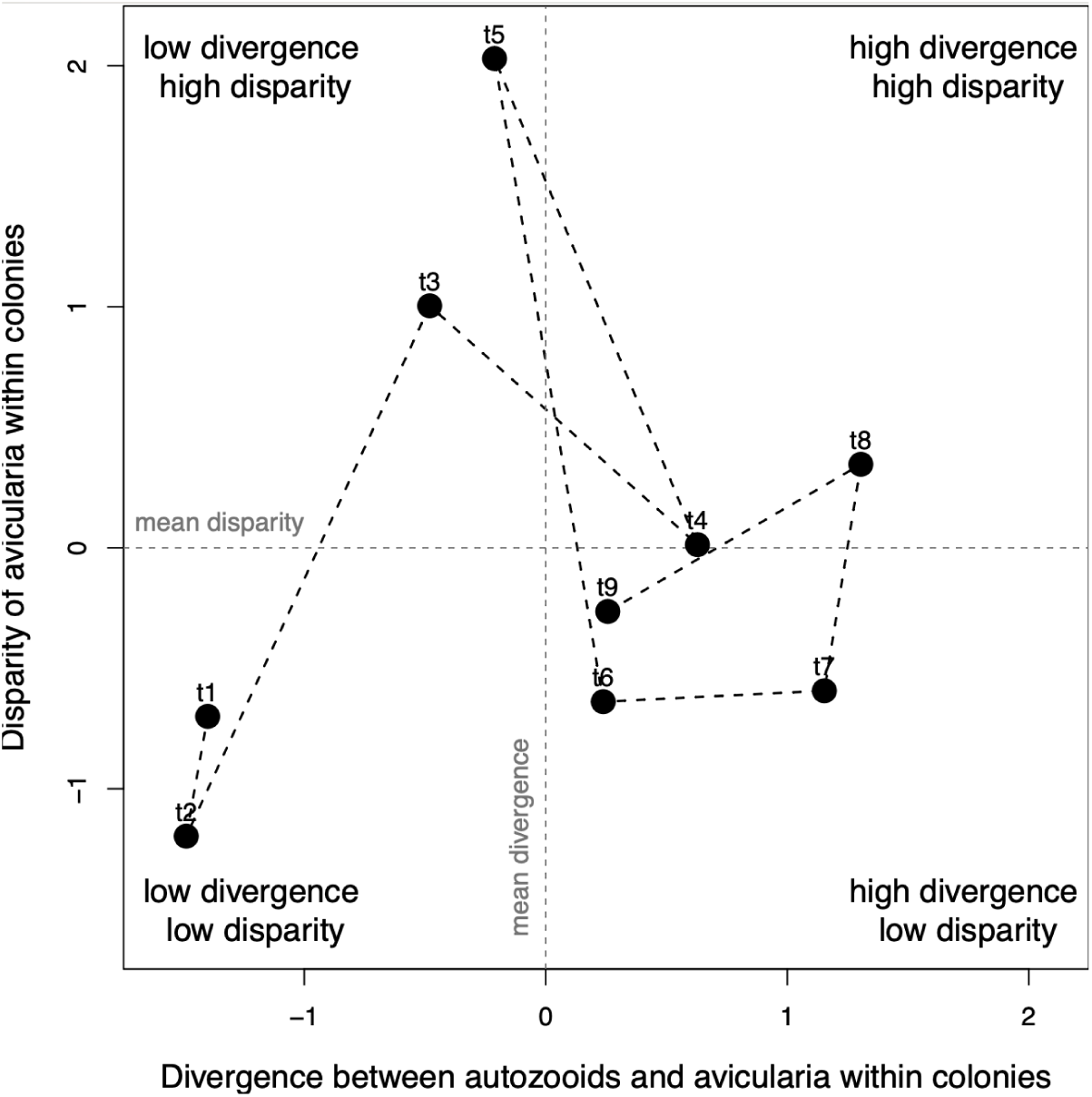
Z-scores of divergence and avicularian disparity within colonies over time. The time series begins with point t_1_ and ends with point t_9_, with each labeled point representing a subsequent formation in the Washita group (see midpoints of time intervals in Fig. 2). Note that the intervals with high avicularian disparity precede intervals with high divergence between autozooids and avicularia within colonies.

The greatest shape divergence between polymorphs occurred within colonies that contained avicularia lacking autozooidal functions, as evidenced from the lack of avicularian ovicells (Fig. 4A, Table S7, following the interpretations of functional avicularian ovicells laid out by Cheetham et al. 2006 (22)). This shape divergence suggests that avicularia, which were derived from autozooids, were apparently released from phenotypic constraints tied to autozooidal functions. The disparity of avicularian shapes tended to be higher within colonies that lacked avicularian ovicells than in those that possessed them (Fig. 4B, Table S8), consistent with the idea that early avicularia could be vestigial autozooids derived from functional release (25, 29–31). Modular developmental decoupling of autozooids and avicularia (32) could then allow selection to act independently on each polymorph, ultimately re-enforcing morphological and functional differences. The loss of ovicells on avicularia in certain colonies also corresponds to a reduction in the overall frequency of ovicells across the colony (Fig. 4A, C, Table S7, Table S9). Decreased frequency of reproductive individuals in a colony appears to be a type of germline sequestration, associated with evolutionary potential at the level of the colony rather than the level of the individual (33, 34). As in many other colonial, social, and multicellular organisms, *Wilbertopora* exhibited a reduction in frequency of reproductive members as new body types emerge (33, 35).

**Figure 4.**
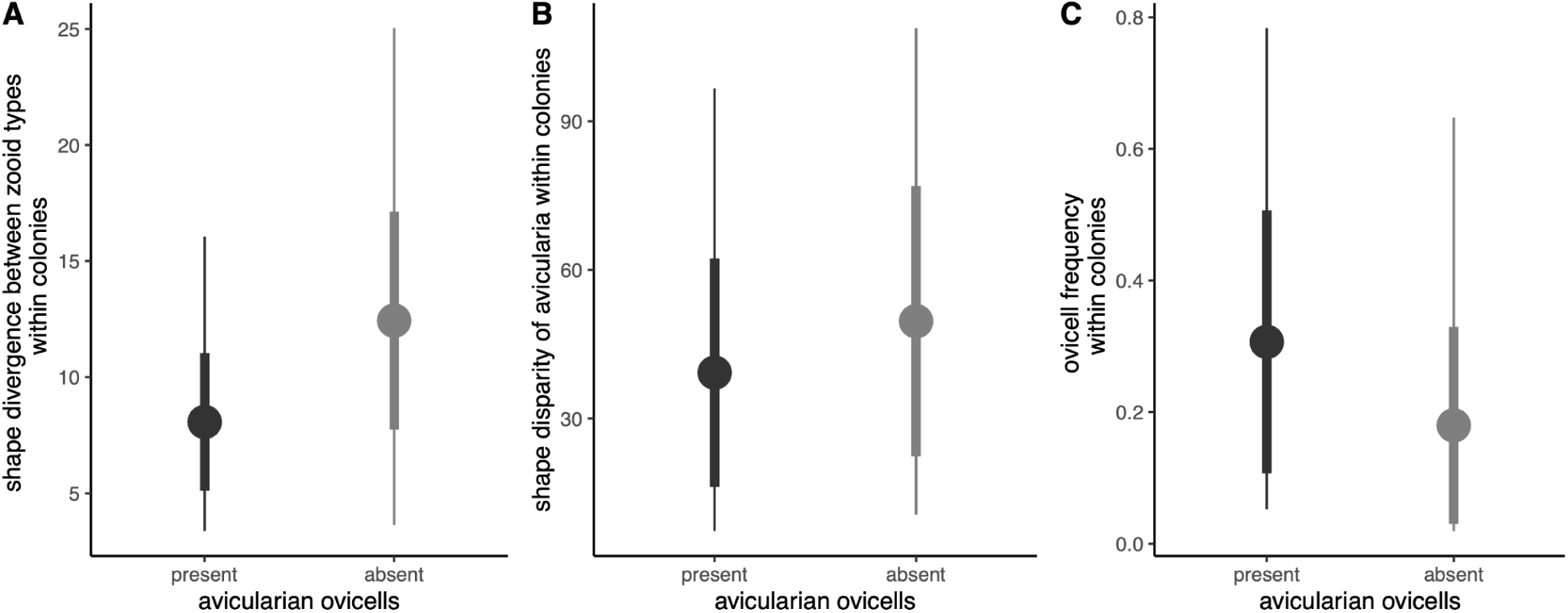
Comparing divergence (**A**), disparity (**B**), and ovicell frequency (**C**) between *Wilbertopora* colonies with and without avicularian ovicells. Points show the mean value, thick line segments show standard deviation, and thin line segments show the range of values. Divergence is significantly different between groups (Table S7, p<0.01), disparity is marginally different between groups (Table S8, p∼0.1), and ovicell frequency is significantly different between groups (Table S9, p<0.01). Results here show colonies with more than 30 zooids and have at least one ovicell.

Taken together, these results show three key features of *Wilbertopora* evolution: (1) avicularia evolved novel morphologies and were not a simple partitioning of pre-existing autozooid morphologies, (2) high pulses of disparity among avicularia suggests that their functional constraints are more relaxed than those of autozooids, and (3) diverging avicularian morphologies suggest potential functional changes from autozooids when they lose associations with ovicells, potentially releasing avicularia to undergo selection for multiple novel functions rather than specializing on pre-existing functions. We propose that the pulses of heightened disparity in avicularian evolution is a transitional phase away from their initially shared functions with autozooids. Gene duplications, as an analog, often result in similar transitional phases where novel variation in one of the gene copies is originally neutral and is later exapted to serve novel functions (36). Early avicularia may have expressed similar neutral variation that lacked functional constraints, but this scenario requires an additional supporting mechanism to explain the persistence of potentially non-functional colony members. Thus, we consider an evolutionary pathway where even nonfunctional avicularia can evolve in colonies despite their apparent energetic cost. This pathway may explain why avicularia have different morphologies among colonies as well as relatively high disparity within colonies.

### Energetic control as a pathway towards division of labor

*Wilbertopora* is one of the first cheilostome bryozoans to exhibit exclusively multiserial growth (3, 16), which brings new geometric arrangements for cheilostome colonies. Prior to *Wilbertopora*, cheilostome lineages had colonies that were primarily uniserial or pluriserial and formed a diffuse network of branches (21), with one exception (37). Zooids in modern uniserial colonies connect to two or three other zooids at most, where old zooids commonly die off after asexual and sexual reproduction and are early colonizers of substrate that soon get overgrown by competitors (16). In contrast, the zooids of *Wilbertopora* colonies form a two-dimensional sheet through their multiserial growth (Fig. 1). Unlike cyclostomes, which possess zooids that can continue to grow via calcification after they asexually bud (38), individual zooids in the interior older parts of a *Wilbertopora* colony are locked into place once a zooid asexually buds and sexually reproduces, and have few further energetic outlets.

Multiserial growth leads to tightly packed zooids, increasing feeding efficiency, and multiplying how much food each zooid acquires compared to each zooid in a uniserial colony (39).

Furthermore, the growing edge of a multiserial colony can be approximated by the circumference of a circle, while zooids comprise its area. Like the ratio of a sphere’s volume to surface area(40), the area of a circle increases faster than its circumference, and this nonlinear relationship has metabolic consequences. As a colony grows, the number of zooids filling its area scales with the colony’s surface area, while the growing edge scales with its circumference. Thus, when all zooids in a colony can feed, the energy required by the growing edge of a colony is exceeded quadratically by the number of feeders in its interior. Nonfeeding zooids can be supported by the colony once it reaches a certain size, and may serve as a sink for otherwise unutilized energetic resources. This would allow colonies to utilize more of the nutrients taken up by autozooids, and would increase the flux of energy through the colony.

We quantified the potential energy flux through a colony by modeling the patterns of nutrient diffusion that are expected to occur given the observed network of zooids (Fig. 5, see supplement). In our model, zooids are networked in what is essentially a bucket brigade, where energy is acquired by feeding zooids, used locally for maintenance, and the excess is passed diffusively from zooid to zooid toward nonfeeding polymorphic zooids and the growing edge, which both act as energy sinks. We ask if the network structure is sufficient to nourish non-feeding members and if the presence of non-feeding members throughout the network increases the energy flux within the colony. As with a bucket brigade, this requires the smooth hand-off of energy from zooid to zooid.

**Figure 5.**
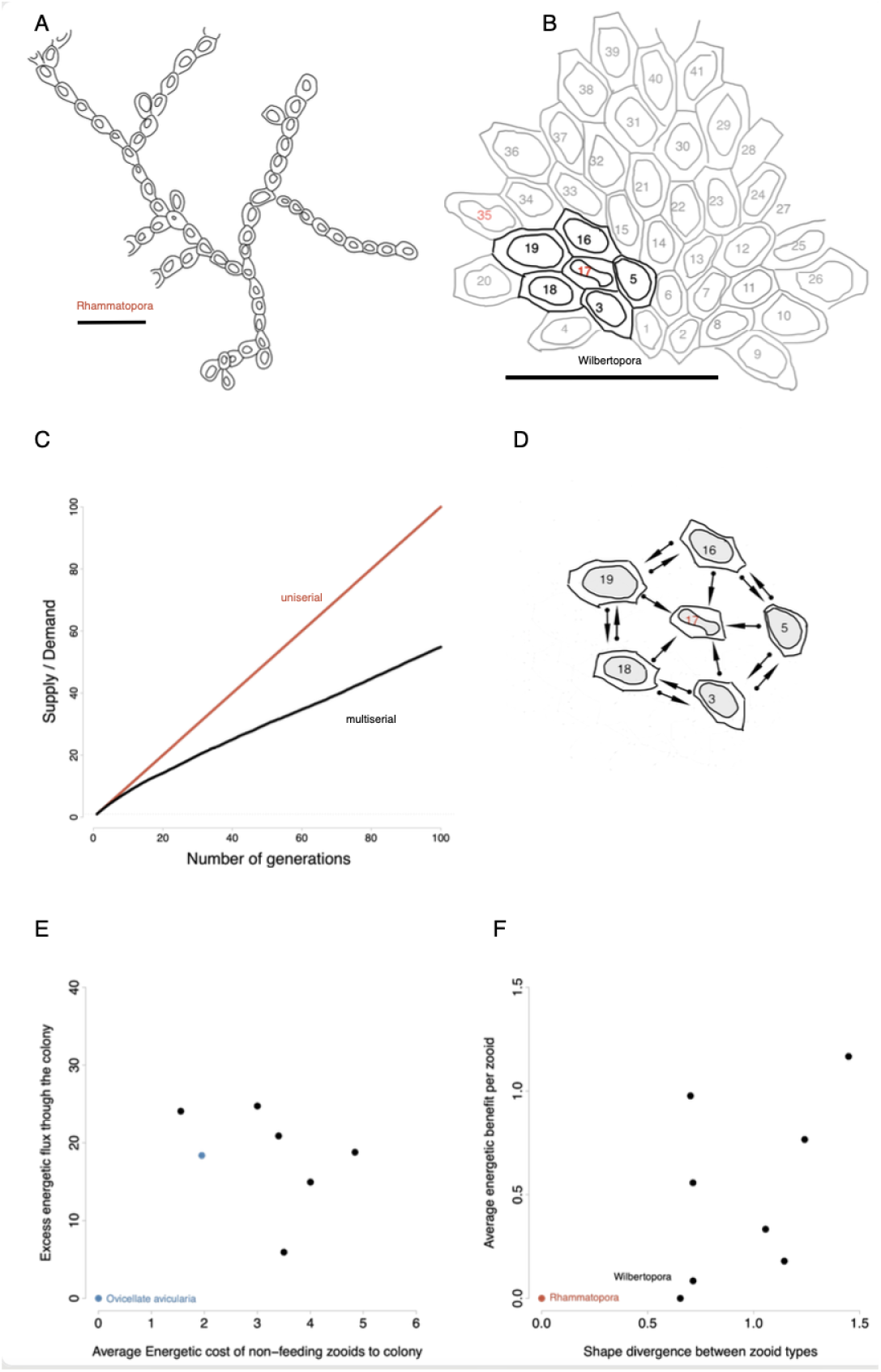
Bryozoan colonies vary in growth form and therefore vary in the connectivity of their physiological connections. We show an illustration of growth forms of (A) the uniserial *Rhammatopora* (PI BZ 8149) and (B) multiserial *Wilbertopora* (USNMPAL 186572). The different budding geometries of each genus lead to different ratios of numbers of feeding zooids to numbers of zooids on the growing edge. (C) Due to physiological connections between zooids, the supply/demand ratio within a colony is a function of the number of feeding zooids and the number of newly budded zooids. Multiserial growth, exhibited in *Wilbertopora*, generates a smaller energy surplus than uniserial growth, indicating higher colony efficiency. The presence of more nonfeeding polymorphs reduces the energy surplus in colonies and works to equilibrate energetic supply with demand. (D) We quantify an incidence matrix of exemplar colonies of *Wilbertopora*. Zooids with red labels in B and D are non-feeding avicularia, zooids with black labels denote feeding autozooids. The funicular system allows the transport of nutrients between zooids. Feeding zooids can transport nutrients between each other, but non-feeding avicularia only receive nutrients from neighbors. Arrows indicate directions of energy exchange. We model diffusion of nutrients along the graph of the colony to evaluate if the colony can support extra non-feeding zooids. We find that adding polymorphic non-feeders within a colony increases the (E) total colony flux of energy and (F) average per zooid energy benefit increases with the presence of non-feeding zooids and their divergence. The contemporaneous genus *Rhammatopora* only has feeding zooids and is plotted as a point of comparison.

We scored incidence matrices (an *n* x *m* matrix with *n* nodes and *m* edges) for exemplar colonies of *Wilbertopora* species, in order to quantify these patterns of energy flow through a colony. We find that the cost of non-feeding members is easily met by pre-existing feeding members, even without serving a function to the colony. As a result, older zooids in the colony serve to increase supply and substantially increase the total energy flux through the colony: up to 100 times the flux per colony (Fig. 5C) and up to 1.25 times the flux per autozooid in a colony (Fig. 5D).

Avicularia can be supported by autozooids that produce more energy than required by the growing edge of the colony and maintenance of existing zooids, even when avicularia serve no distinct functional role. Such lack of function serves to disconnect the origins of division of labor from polymorphism and introduces a phase where the loss of function can precede any gain of function. The energetic surplus due to multiserial growth releases non-feeding members from functional constraints and permits them to vary, much as one copy of a duplicated gene is free to vary. After species pass through this phase of functional loss, evidenced by the decline in avicularia having ovicells, we observe distinct regions of morphospace occupied by different species of *Wilbertopora* (Fig. 2 A). The avicularian phenotypes of those species are diverse and already resemble phenotypes similar to those that occur in modern avicularia with distinct functions(22, 41). As with gene duplication, independent lineages may follow different evolutionary paths. This result explains why it is so difficult to pinpoint a single ancestral function of avicularia, because the later species of *Wilbertopora* already possess avicularian forms similar to the general classes of avicularian types observed today (14).

Energy sinks also promote increased uptake of nutrients from the water column, which along with increased feeding efficiency (39), may serve to deplete food resources for competing encrusting animals (42, 43). Furthermore, this energetic surplus may explain how brooding behavior, a costly reproductive strategy(44), has become the dominant mode of reproduction in cheilostomes, as well as how subsequent evolutionary novelties in cheilostomes, such as multizooidal budding and frontal avicularia, evolve. These energetic aspects of cheilostomes’ ecological success may explain their rapid diversification through much of the Cretaceous and Cenozoic(19, 20). This adds energy flux to their other known ecological strategies of dominant cheilostomes including flow patterns (45), feeding efficiency(39), and budding patterns (16).

### Conclusions

Colonial organisms that consist of only one member type have a life-history that is equivalent to the life-histories of its members; polymorphism allows colonies to evolve novel life-history strategies that vary as a function of the frequencies of their polymorph types(46). Beyond the combinatorics of the member types, we find that polymorphs may not be so costly because they add an inherent energy flux advantage. Together, this expansion of the possible life-history strategies may contribute to heightened speciation rates and extinction rates as the complex competitive networks grow, including non-transitive interactions (47) among cheilostomes.

Polymorphism has long held an air of mystery. It is simultaneously associated with highly successful clades of colonial organisms, but it is also relatively rare across taxa(33). Because of how important economic ideas of specialization have been(48) in understanding division of labor, and the evolutionary idea that the functional loss, such as altruism, is difficult to evolve (2, 49, 50), equating morphological polymorphism with division of labor has led to little progress in understanding either. Darwin’s discussion of polymorphism in bryozoans (51) concerns the functional gains at the origin of polymorphism and reads as remarkably modern (41) because we have made so little progress in our understanding. The problem equating polymorphism with economic division of labor is that it hides how functions can evolve(52) and hides how the developmental origins of novel features are distinct from their role in ecological innovation(53). We find that phenotypic polymorphism within colonies is best thought of as any other biological novelty: how new forms developmentally arise and what ecological advantages they initially may, or may not, provide.

## Methods

### Specimen sampling

All specimens used in this study are housed at the National Museum of Natural History (NMNH) in Washington, D.C. Specimens were collected primarily between 1937 and 1946, with some additional specimens collected in the 1990s, from outcrops of the Washita Group located in North-central Texas (54–56). For a full description of specimen collection from the field, see (57).

While species identification was recorded for the dataset, our analysis considers colony-level, and not species-level, variation, so all formations in the Washita group can be considered as a time series. Species are largely defined by their avicularian morphology, making it difficult to independently consider cladistic evolution and morphological differentiation of avicularia.

Additionally, Cheetham et al (2006)(22) hypothesized that there may be ghost lineages in *Wilbertopora*. This makes it difficult to estimate rates of trait evolution. However, for this study, we focus on trait differences between geologic formations rather than rates of evolutionary change. Furthermore, we use more colonies than Cheetham et al (2006) included in their analysis, and all species considered in this analysis are verifiably descended from a recent common ancestor (22, 58, 59). More information of species categorization can be found in the Supplement sections 1 and 5. Museum registration numbers can be found in Dataset S1.

### Quantification and statistical analysis

#### Digitization and quantification of zooid orifice and opesia morphology

Bryozoan colonies grow on irregular surfaces, making repeatable capture of the orifice and opesia shape in the orificial plane difficult. Therefore, two approaches were taken to minimize the effect of parallax on the orifice and opesia shape: micro-CT scanning to create 3D models for colonies growing on curved surfaces, and microscope photography of colonies growing on flat surfaces. Specific descriptions of the steps for followed for the acquisition of both micro-CT scans and photographs can be found in the Supplement section 2. Landmarks were placed on mesh models using ‘Pick Points’ in Meshlab (60). Landmarks were placed on photographs using FIJI (61), and scaled from pixels to millimeters to match units of mesh data.

Outlines from mesh data were projected onto the orificial plane to match the outlines derived from photographs. A two-dimensional spline was used to place 50 equally spaced semilandmarks along the curve defined by the initial outline points. Semilandmarks were then slid to minimize bending energy and reduce artifactual differences in shape driven by their initial, equidistant placement(62–64). The semilandmark configurations were centered on their respective centroids and aligned using the distal and proximal landmarks. The dimensionality of these aligned semilandmark configurations was reduced using principal components analysis (PCA, Fig. S2). The differences in avicularia and autozooid morphology among colonies through time was tested via a residual randomization permutation procedure (using RRPP::lm.rrpp (65)) on a linear model of PC scores against zooid type interacting with time (midpoint in Mya per formation).

### Estimation of morphological divergence between zooid types

To test for differences in the shapes of zooid types through time, Euclidean distances were calculated between the predicted mean autozooid shape in the first time interval to the predicted mean shapes for each zooid types per time interval across the 999 coefficients derived via the RRPP procedure (uncertainty expressed as the 95% confidence interval). The differences in avicularia and autozooid morphology within colonies through time was tested by first finding the Euclidean distance between the mean shapes of zooid types within colonies (distances based on mean and median morphologies were similar, Fig. S4), and then regressing that distance against time (model 4, Table S4, Fig. S5).

### Estimation of disparity

Shape disparity for each zooid type across colonies was estimated using geomorph::morphol.disparity(62), and temporal trends were evaluated by regressing disparity across each time interval using RRPP:lm.rrpp (S7, S5). The regression model (model 6) tests for differences of colony-level disparity across zooid types and time intervals. The resulting predicted values for disparity show that, while avicularia have more disparity than autozooids in colonies, there is no monotonic temporal trend for disparity (Fig. S8, Table S6).

### Avicularian ovicells and their relationship to divergence and disparity

Ovicells that grow on avicularia suggest that some avicularia retain ancestral autozooidal functions. For specific discussion of the significance of avicularian ovicells, see Supplement section 7. Colonies with ovicells were checked for the presence of ovicells on avicularia using meshes and photographs. Colonies that lack ovicells or fall below a certain size (<30 zooids) were excluded from this portion of the analysis because smaller colonies may not have reached sexual maturity yet. For each colony, all zooids were counted to estimate colony size, and ovicells were counted to estimate ovicell frequency.

Linear modeling was used to determine whether there is a significant relationship between divergence within colonies and the presence or absence of avicularian ovicells in colonies (model 7, Table S7). Linear modeling was also used to determine whether there is a significant relationship between avicularian disparity within colonies and the presence of avicularian ovicells within colonies (model 8, Table S8). To estimate whether ovicell frequency differs between colonies that possess ovicells on avicularia and those that lack them, ovicell frequency was regressed against the status of avicularian ovicells (present, absent) in colonies (model 9, Table S9).

### Additional resources

We model energy flux using an incidence matrix where each zooid in a colony is a node. Model code and explanation are included in supplemental materials (Section 8).

## Acknowledgements

We thank J. Sanner and J.B.C. Jackson for their collection and maintenance of the bryozoan collection. We thank D. Erwin, G. Hunt, and M. Fau for discussion of the manuscript. This research was supported by the Smithsonian Institution Fellowship Program and the Paleontological Society student research grant. All data and materials are available in the supplementary materials.

## Author contributions

S.L., S.M.E., and C.S. developed the project idea, S.L. collected the data, S.L., S.M.E., C.S., and R.M. conducted the data analysis, S.L., S.M.E., and C.S. wrote and edited the manuscript.

## List of Supplementary Materials

Supplementary Text

Figs. S1 to S9

Tables S1 - S9

Data files S1-S3

