## Supplemental materials for "Origin of division of labor is decoupled from polymorphism in colonial animals"

### 1 Specimen Sampling

All specimens used in this study are housed at the National Museum of Natural History (NMNH) in Washington, D.C. Specimens were collected primarily between 1937 and 1946, with some additional specimens collected in the 1990s, from outcrops of the Washita Group located in North-central Texas (1, 2, 3). For a full description of specimen collection from the field, see Cheetham 1954 and Cheetham et al. 2006 (4, 5).

The Washita Group is composed of nine sequential formations (Kiamichi, Duck Creek, Fort Worth, Denton, Weno, Paw Paw, Main Street, Georgetown, Grayson Formations), spanning approximately 9 Ma (∼103 Ma to 96 Ma) (1, 3, 6). The age ranges for each formation that are used in this study are derived from chronostratigraphic studies by Scott et al (3, 6).

Many of these specimens were assigned to a specific species of *Wilbertopora* (5), but most of the specimens found in the beginning and middle portions of the Washita Group (Kiamichi Formation, Duck Creek Formation, Weno Formation, Paw Paw Formation) lack species identifications, due to morphology that is inconsistent with named species in (5).While species identification was recorded for the dataset, our analysis primarily considers colony-level, and not species-level, variation, so all formations in the Washita group can be considered as a time series. Species are largely defined by their avicularian morphology, making it difficult to independently consider cladistic evolution and morphological differentiation of avicularia. Additionally, Cheetham et al (2006) (5) hypothesized that there may be ghost lineages in Wilbertopora. This makes it difficult to estimate rates of trait evolution. However, for this study, we focus on trait differences between geologic formations rather than rates of evolutionary change. Furthermore, we use more colonies than Cheetham et al (2006) included in their analysis, and all species considered in this analysis are verifiably descended from a recent common ancestor (5, 7, 8). More information of species categorization can be found in the Supplement section 5. Museum registration numbers can be found in Dataset S1.

Specimens with poor preservation of zooid frontal area morphology were excluded from the study.

Museum registration numbers can be found in Dataset S1.

### 2 Digitization of zooid frontal area

#### 2.1 Specimen capture

Bryozoan colonies grow on irregular surfaces, making repeatable capture of the orifice and opesia (together comprising the frontal area) shape in the frontal plane difficult. Therefore, two approaches were taken to minimize the effect of parallax on the orifice and opesia shape:

micro-CT scanning to create 3D models for colonies growing on curved surfaces, and microscope photography of colonies growing on flat surfaces.

**Micro-CT scanning.** Specimens were scanned using micro-computed X-ray tomography ($\mu$CT) at the National Museum of Natural History Scientific Imaging Facility. Three-dimensional, isosurface, triangular-mesh models were created in VG Studio Max and ORS Dragonfly. All meshes are available online via links in electronic supplementary material, Dataset S1.

**Photography.** For specimens that had features not resolved at practical microCT resolution (˜10-15µm), or for specimens that are encrusted on surfaces too large for microCT scanning, microscope photography was used to capture morphological features. Only specimens on flat surfaces were considered for microscopy, so the parallax effect could be minimized. Specimens were photographed in single focal planes or small z-stacks (<5 photos per stack) to ensure minimal distortion of zooid features. Photographs were taken using a Nikon DS-Fi3 and photographs were prepared using Nikon elements software.

#### 2.2 Frontal area outlines

A zooid's frontal area is the main opening of its body cavity, where the soft tissue (polypide) of the animal is found in life, and from where the polypide can be extruded for feeding (9). The orifice is met by the opesia at its distal end, and together these features make up the frontal area of a zooid, covered by the frontal membrane in life.The orifice and opesia shape is defined by the edge of the gymnocyst, which is the calcified outer wall of the zooid body. Each zooid's frontal area was initially digitized with sufficient density to capture its shape, beginning at the most distal end of the zooid and moving clockwise around the orifice and opesia. Landmarks were placed at the most distal and proximal portions of the zooid orifice and opesia, so outlines could be transformed to have identical orientations. Landmarks were placed on mesh models using 'Pick Points' in Meshlab (10). Landmarks were placed on photographs using FIJI (11), and scaled from pixels to millimeters to match units of mesh data.

### 3 Quantifying frontal area morphology

Outlines from mesh data were projected onto the frontal plane to match the outlines derived from photographs. A two-dimensional spline was used to place 50 equally spaced semilandmarks along the curve defined by the initial outline points. The semilandmark on the curve nearest the distal landmark (as defined above) was selected as the initial point for the curve (Fig. S1). Semilandmarks were then slid to minimize bending energy and reduce artifactual differences in shape driven by their initial, equidistant placement (12, 13) using the R function *geomorph::gpagen* (14). The semilandmark configurations were centered on their respective centroids and aligned using the distal and proximal landmarks. The dimensionality of these aligned semilandmark configurations was reduced using principal components analysis (PCA, Fig. S2). PCs 1-9 (explaining 95% of the total variation in shape) were retained for analyses of orifice shape.

Orifice and opesia size was measured as the centroid size of each orifice and opesia outline.

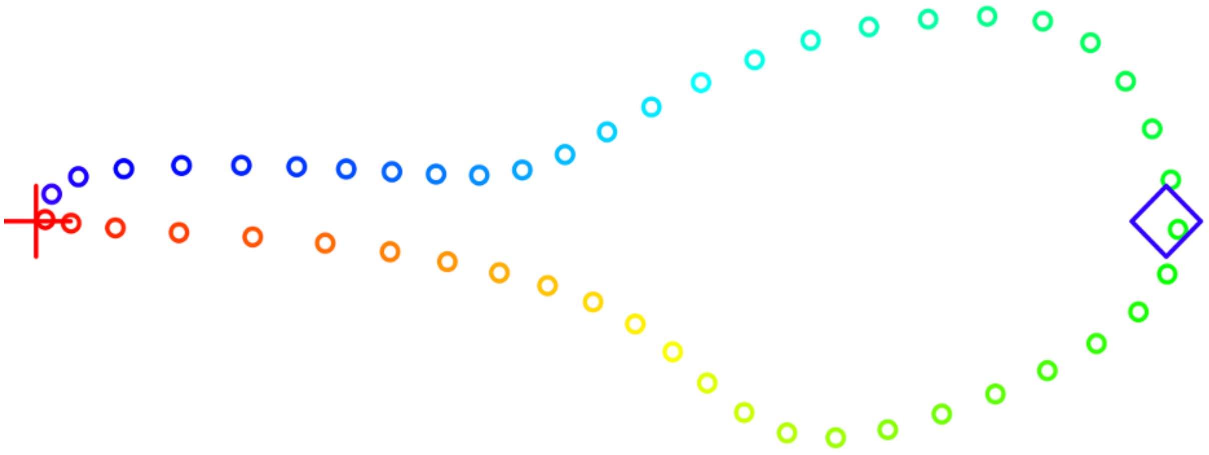

Figure S1: Example outline of avicularium after spline-fit and slid semi-landmarks. Outline starts with red points and ends with blue points. Distal landmark as red cross, proximal landmark as blue diamond.

### 4 Relationship between size and shape of zooid orifices and opesia

The R package RRPP (*RRPP::lm.rrpp*) was used for linear modeling. To determine whether shape and size are independent variables, shape was regressed against size:

**Model 1:** $lm.rrpp(PC_{1-9} \sim CS)$

Shape is significantly correlated with size, but with high unexplained variance (Table S1).

To test whether size-shape relationships vary across zooid types, shape was regressed onto size and zooid type: **Model 2:** $lm.rrpp(PC_{1-9} \sim CS * type)$

Shape is significantly different between zooid types while accounting for any differences in their size (Table S2), but shape does not interact with size and zooid type. Thus, autozooids

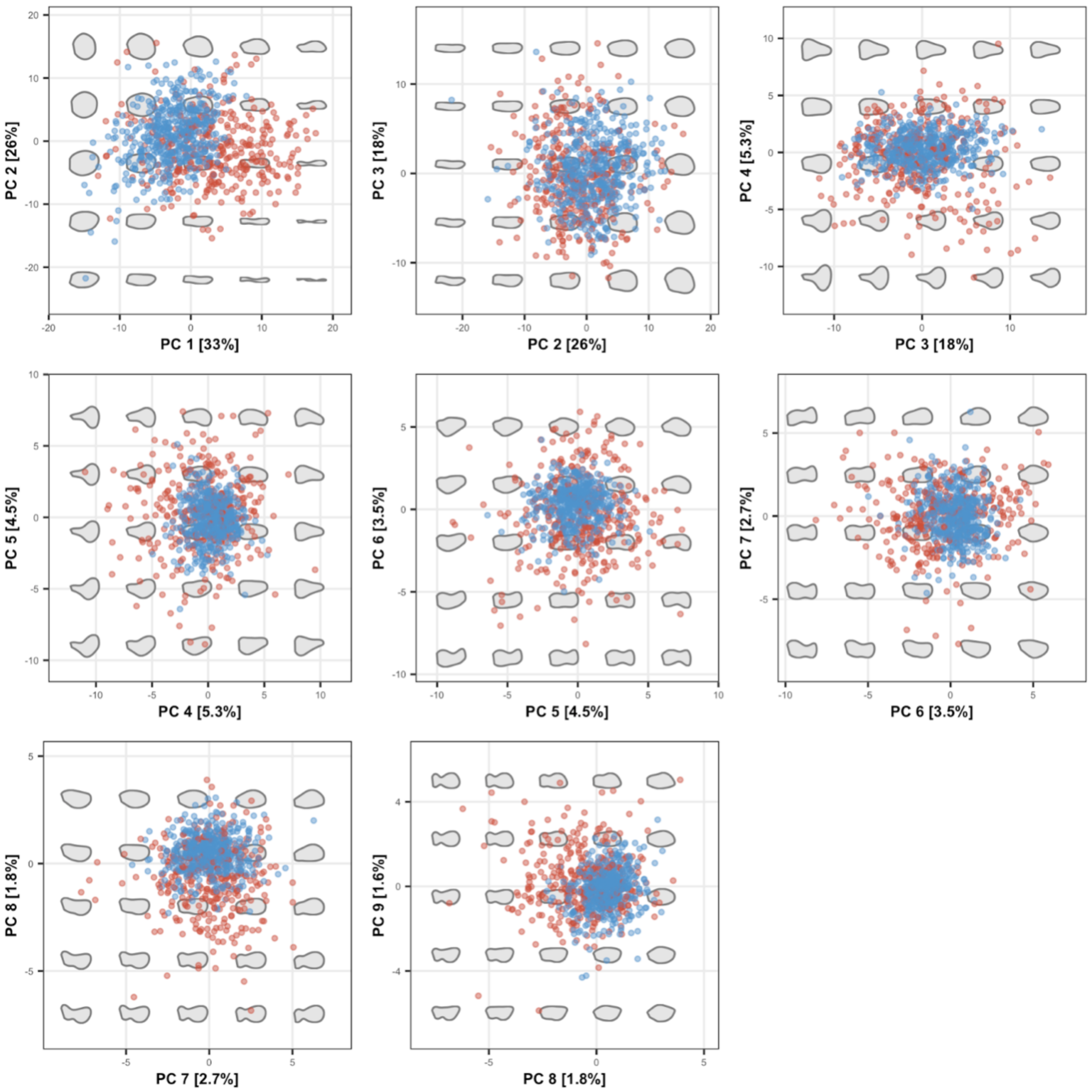

Figure S2: Morphospace of zooid orifices along the first nine PCs. Avicularia as red points, autozoids as blue points. Shapes are the latent projections of orifice shape at specified PC scores.

Table S1: Analysis of variance (ANOVA) for regression of orifice and opesia shape as a function of zooid centroid size (model 1). Terms=effects in ANOVA, Df=degrees of freedom, SS=Sum of Squares, MS=Mean Squares.

| **Terms** | **Df** | **SS** | **MS** | **$R^2$** | **F** | **Z** | **Pr>F** |
|---|---|---|---|---|---|---|---|
| *CS* | 1 | 4190 | 4190 | 0.0532 | 46.1 | 8.04 | 0.001 |
| Residuals | 822 | 74600 | 90.7 | 0.947 | | | |
| Total | 823 | 78800 | | | | | |

and avicularia are offset in their size-shape correlations, but this offset remains constant over the range of observed zooid sizes. Given these correlations, results are interpreted with respect to shape differences, which reflect functional differences in polymorphs of extant bryozoans (9, 15).

Table S2: Analysis of variance (ANOVA) for regression of orifice and opesia shape as a function of zooid type interacting with zooid centroid size (model 2). Table header as in Table S1.

| **Terms** | **Df** | **SS** | **MS** | **$R^2$** | **F** | **Z** | **Pr>F** |
|---|---|---|---|---|---|---|---|
| *CS* | 1 | 4190 | 4190 | 0.0532 | 52.7 | 8.37 | 0.001 |
| *type* | 1 | 9410 | 9410 | 0.119 | 119 | 8.12 | 0.001 |
| *CS:type* | 1 | 75.7 | 75.7 | $9.61*10^{-4}$ | 0.954 | 0.954 | 0.41 |
| Residuals | 820 | 65100 | 79.4 | 0.826 | | | |
| Total | 823 | 78800 | | | | | |

### 5 Shape Divergence

#### 5.1 Divergence across colonies

The differences in avicularia and autozooid morphology *among* colonies through time was tested via a residual randomization permutation procedure (using *RRPP::lm.rrpp* (16)) on a linear model of PC scores against zooid type interacting with time (midpoint in Mya per formation):

**Model 3:** $lm.rrpp(PC_{1-9} \sim type + time + type : time)$

Autozooid and avicularia morphology significantly diverge in shape through time (Table S3), the marginal effect of *time* is significant after accounting for differences in *type* and that shape depends on both *type* and *time*).

Table S3: Analysis of variance (ANOVA) for regression of orifice and opesia shape as a function of zooid type interacting with time (model 3). Table header as in Table S1.

| **Terms** | **Df** | **SS** | **MS** | **$R^2$** | **F** | **Z** | **Pr>F** |
|---|---|---|---|---|---|---|---|
| *type* | 1 | 1671 | 1671 | 0.022 | 20 | 6.2 | 0.001 |
| *time* | 1 | 1684 | 1684 | 0.023 | 20 | 6.1 | 0.001 |
| *type:time* | 1 | 1540 | 1540 | 0.021 | 19 | 6.1 | 0.001 |
| Residuals | 776 | 64069 | 82 | 0.86 | | | |
| Total | 779 | 74477 | | | | | |

To test for differences in the shapes of zooid types through time, Euclidean distances were calculated between the predicted mean autozooid shape in the first time interval (i.e. its estimated coefficient from model 3) to the predicted mean shapes for each zooid types per time interval across the 999 coefficients derived via the RRPP procedure (uncertainty expressed as the 95% confidence interval). Using this approach, both autozooids and avicularia shapes showed significantly increased distances to the autozooid morphology from the first time interval (Fig. S3), with avicularia showing greater shape differences through time.

#### 5.2 Divergence within colonies

The differences in avicularia and autozooid morphology *within* colonies through time was tested by first finding the Euclidean distance between the mean shapes of zooid types within colonies ($DST_{\mu}$; distances based on mean and median morphologies were similar, Fig. S4), and then regressing that distance against time:

**Model 4:** $lm.rrpp(DST_{mu} \sim time)$

Autozooid and avicularia morphology significantly diverge in shape through time within colonies (Table S4, Fig. S5).

#### 5.3 Operational species-level divergence within colonies

To estimate how species-level dynamics may underpin broader patterns of divergence *Wilbertopora* colonies, divergence values were grouped by species (Fig. S6). Due to nonindependence of orifice and opesia shape and size traits with species assignments, it is expected that different species would express different ranges of divergence.

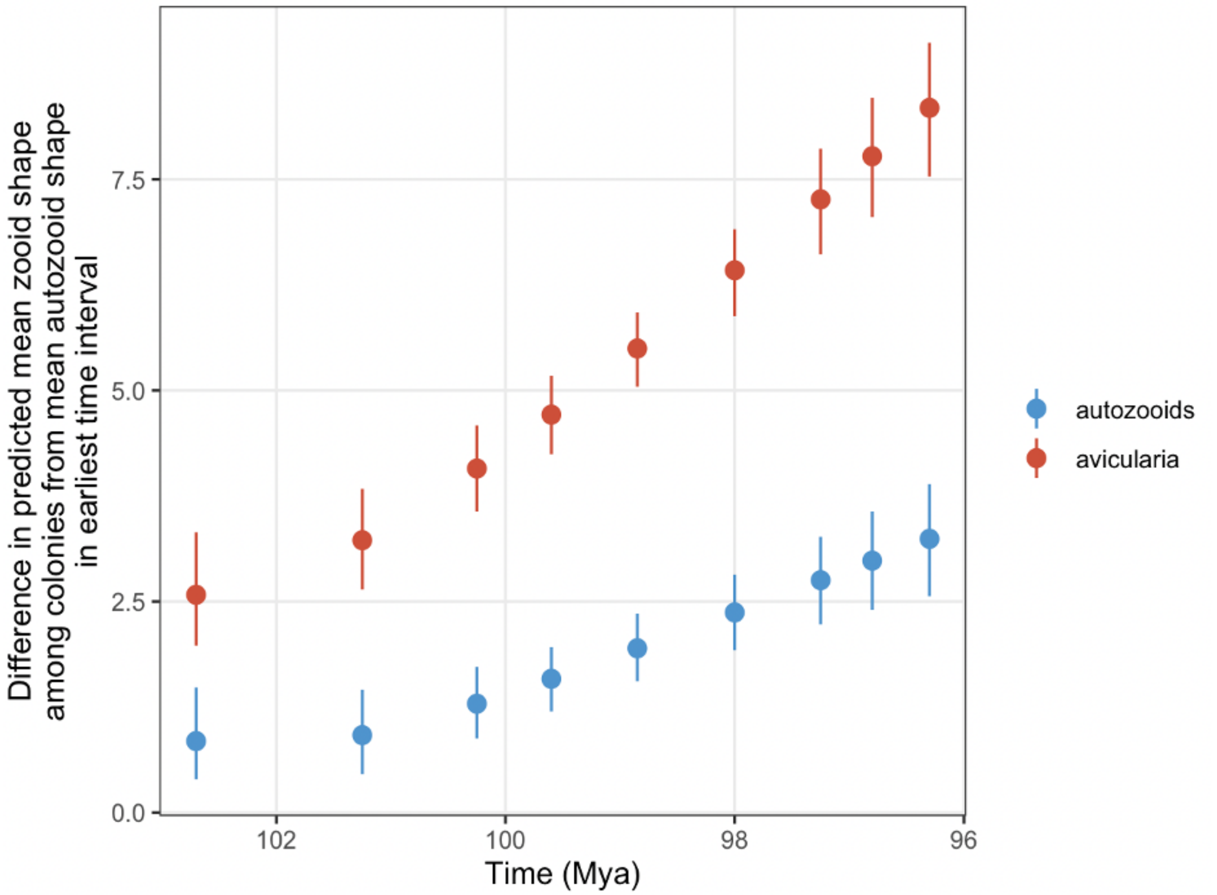

Figure S3: Euclidean distances between predicted mean shape for autozooids in the first time interval and the predicted mean shape for autozooids (blue) and avicularia (red) in each interval. 95% confidence intervals are shown as bars around each point estimate.

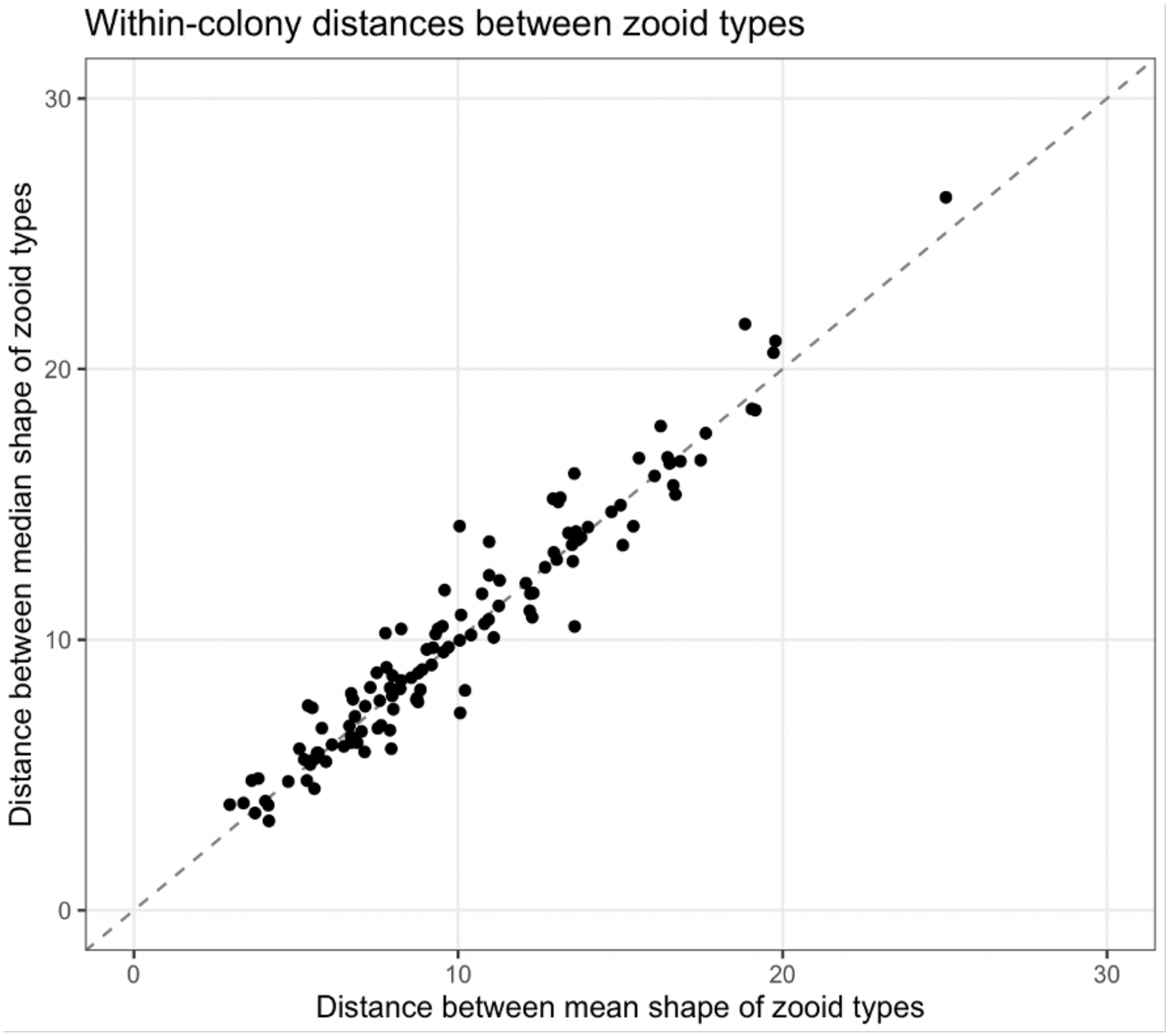

Figure S4: Comparison of Euclidean distances between mean and median shapes of zooid types in colonies. Each point is a colony; dashed gray line shows 1:1.

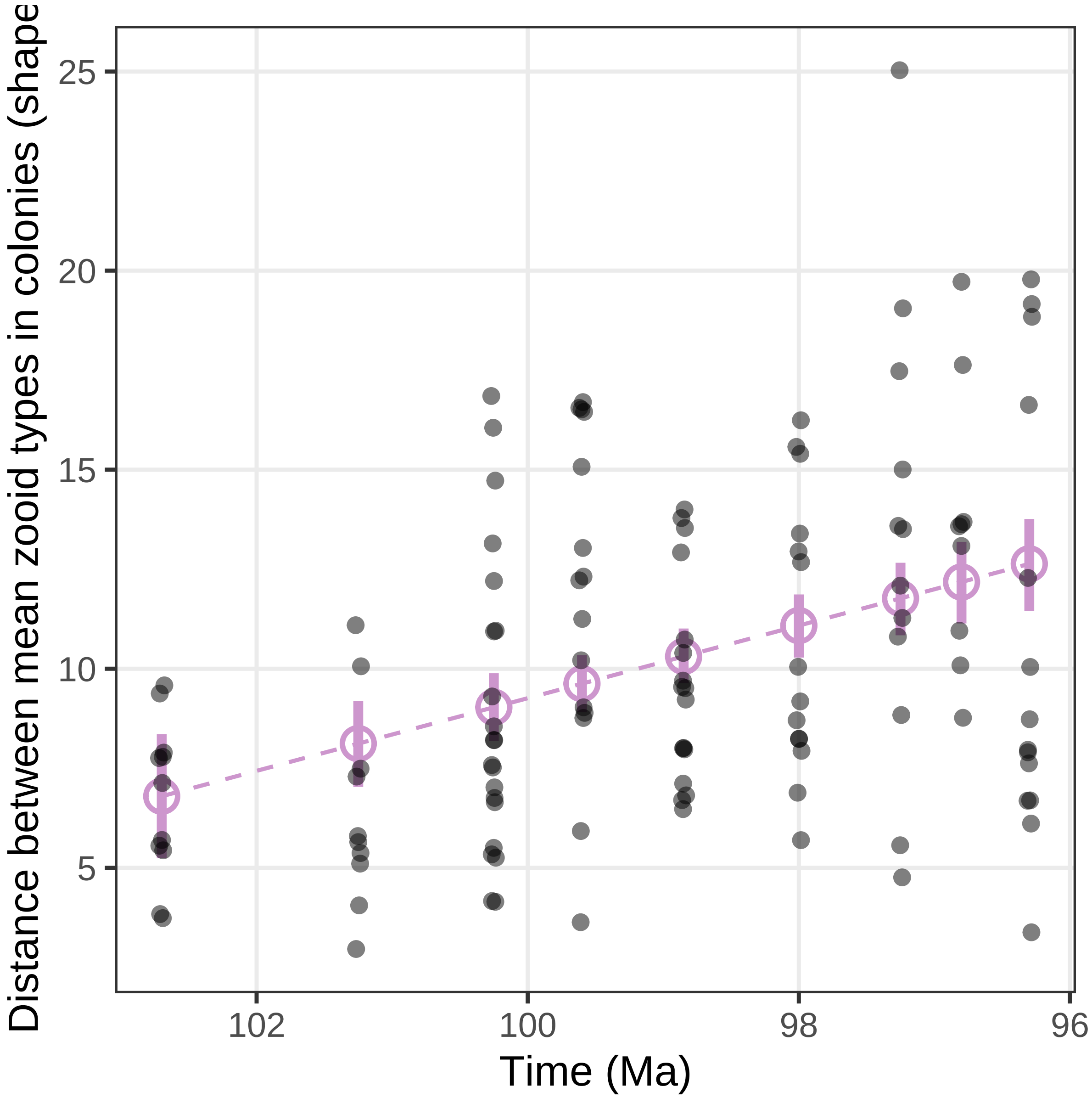

Figure S5: Distance between mean orifice and opesia *shape* morphology for each zooid type in each colony. Black points represent the distance value for each colony, and the light purple points represent predicted mean values based on model 4. Lines around each mean predicted value represent 95% confidence intervals.

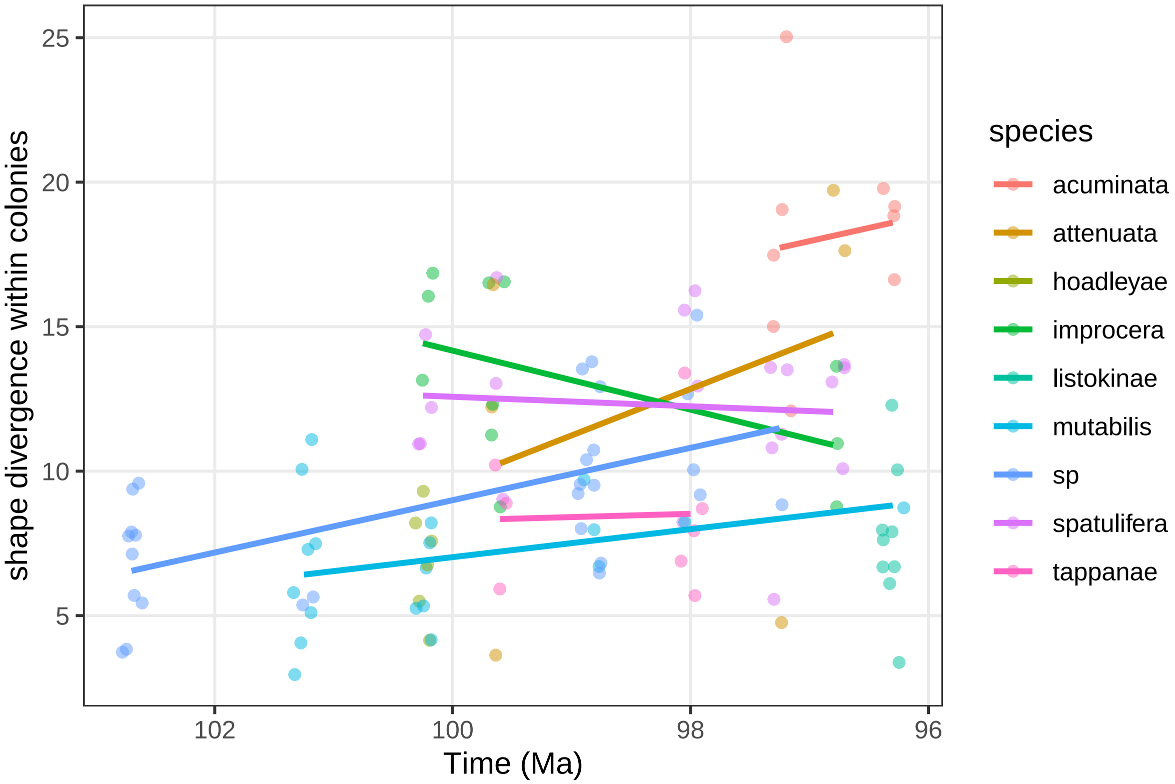

Figure S6: Divergence within colonies over each time interval and across species.

Table S4: ANOVA for regression of Euclidean distances between mean orifice and opesia shapes of zooid types within colonies as a function of time (model 4). Table header as in Table S1.

| **Terms** | **Df** | **SS** | **MS** | $\mathbf{R^2}$ | **F** | **Z** | **Pr>F** |
|---|---|---|---|---|---|---|---|
| *time* | 1 | 365 | 365 | 0.161 | 23.1 | 3.52 | 0.001 |
| Residuals | 121 | 1910 | 15.8 | 0.839 | | | |
| Total | 122 | 2280 | | | | | |

### 6 Estimation of disparity

#### 6.1 Shape disparity across colonies

Shape disparity for each zooid type *across* colonies was estimated using *geomorph::morphol.disparity*, and temporal trends were evaluated by regressing disparity across each time interval using *RRPP:lm.rrpp*:

$$Disp < -\, morphol.disparity(PC_{1-9} \sim time * type, groups = interaction(time, type))$$

**Model 5:** $lm.rrpp(Disp \sim time * type)$

While disparity across colonies significantly differs between zooid types, there is no significant trend across time (S7, S5). The lack of significant change between zooid types over time is likely a result of large uncertainties around each point estimate, which were derived from the predicted disparity values of the model (model 5). Nevertheless, avicularian orifices and opesia have significantly more shape disparity than autozooids, reflecting their expansion into novel regions of morphospace.

Table S5: ANOVA for regression of distances between mean orifice and opesia shapes of zooid types within colonies as a function of time (model 5). Table header as in Table S1.

| **Terms** | **Df** | **SS** | **MS** | $\mathbf{R^2}$ | **F** | **Z** | **Pr>F** |
|---|---|---|---|---|---|---|---|
| *type* | 1 | 9320 | 9320 | 0.588 | 24.3 | 3.2 | 0.001 |
| *time* | 1 | 582 | 582 | 0.0367 | 1.52 | 0.691 | 0.258 |
| *type:time* | 1 | 573 | 573 | 0.588 | 1.49 | 0.696 | 0.248 |
| Residuals | 14 | 5370 | 383 | 0.339 | | | |
| Total | 17 | 15800 | | | | | |

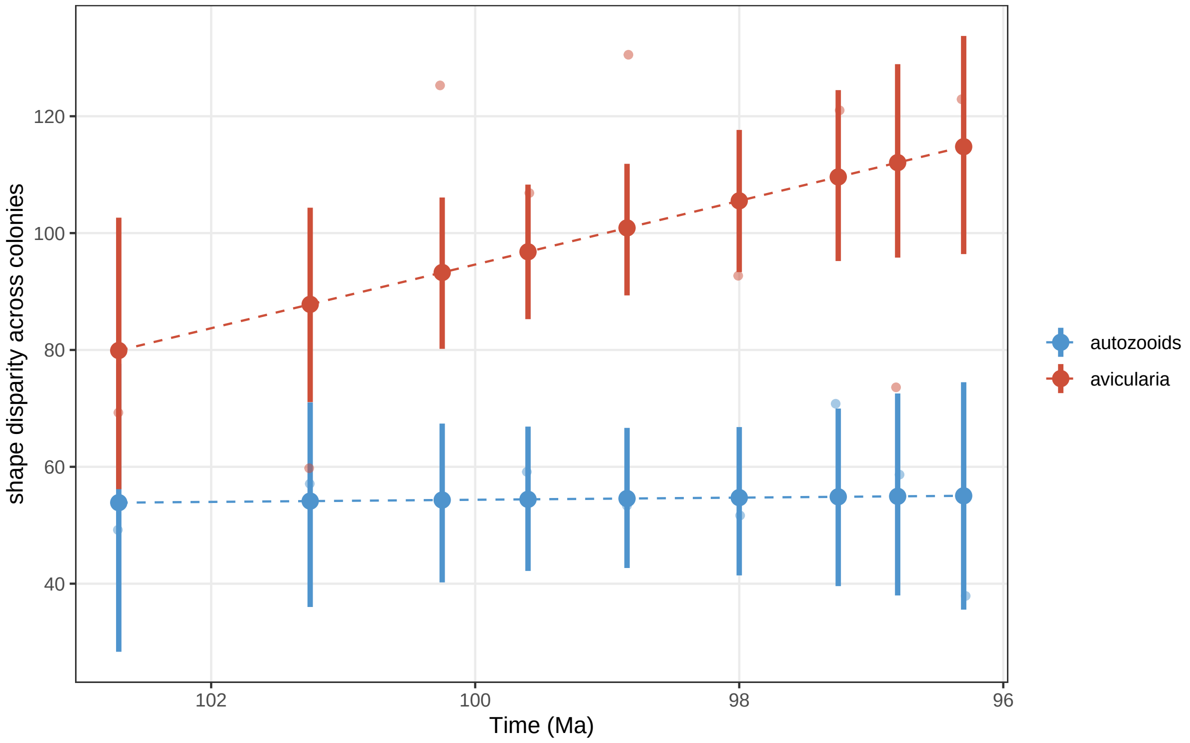

Figure S7: Shape disparity across colonies over time. Disparity values for each zooid type in each time interval are shown as transparent points. Mean predicted disparity values are shown as opaque points, and 95% confidence intervals are shown as segments around the points. On average, avicularia have higher disparity than autozoids across colonies through time, but the apparent interaction between zooid type and shape disparity is not significant (Table S5).

#### 6.2 Shape disparity within colonies

The disparity of orifice and opesia shape *within* colonies was estimated similarly as in model 5, but with an added interaction of colony identity:

$Disp <- morphol.disparity(PC_{1-9} \sim colony * time * type,$
$groups = interaction(colony, time, type))$

**Model 6:** $lm.rrpp(Disp \sim time * type)$

The regression model (model 6) tests for differences of colony-level disparity across zooid types and time intervals. The resulting predicted values for *Disp* show that, while avicularia have more disparity than autozooids in colonies, there is no temporal trend for disparity (Fig. S8, Table S6). This result suggests that autozooids and avicularia maintain their levels of disparity in colonies over the course of *Wilbertopora*'s evolution in the Washita Group.

Table S6: ANOVA for regression of distances between mean orifice opesia shapes of zooid types within colonies as a function of time (model 6). Table header as in Table S1.

| **Terms** | **Df** | **SS** | **MS** | **$R^2$** | **F** | **Z** | **Pr>F** |
|---|---|---|---|---|---|---|---|
| *type* | 1 | 15200 | 15200 | 0.0761 | 20 | 3.37 | 0.001 |
| *time* | 1 | 35 | 35 | $1.75*10^{-4}$ | 0.046 | -1.09 | 0.851 |
| *type:time* | 1 | 256 | 256 | 0.00128 | 0.337 | -0.147 | 0.576 |
| Residuals | 242 | 184000 | 762 | 0.922 | | | |
| Total | 245 | 200000 | | | | | |

#### 6.3 Operational species-level shape disparity within colonies

To estimate how species-level dynamics may underpin broader patterns of disparity in *Wilbertopora* colonies, disparity values were grouped by species (Fig. S9).

### 7 Avicularian ovicells

#### 7.1 Theoretical background

Connecting the morphological differentiation of polymorphs to potential functional differentiation is difficult without the existence of extant representatives of *Wilbertopora*. Moreover,

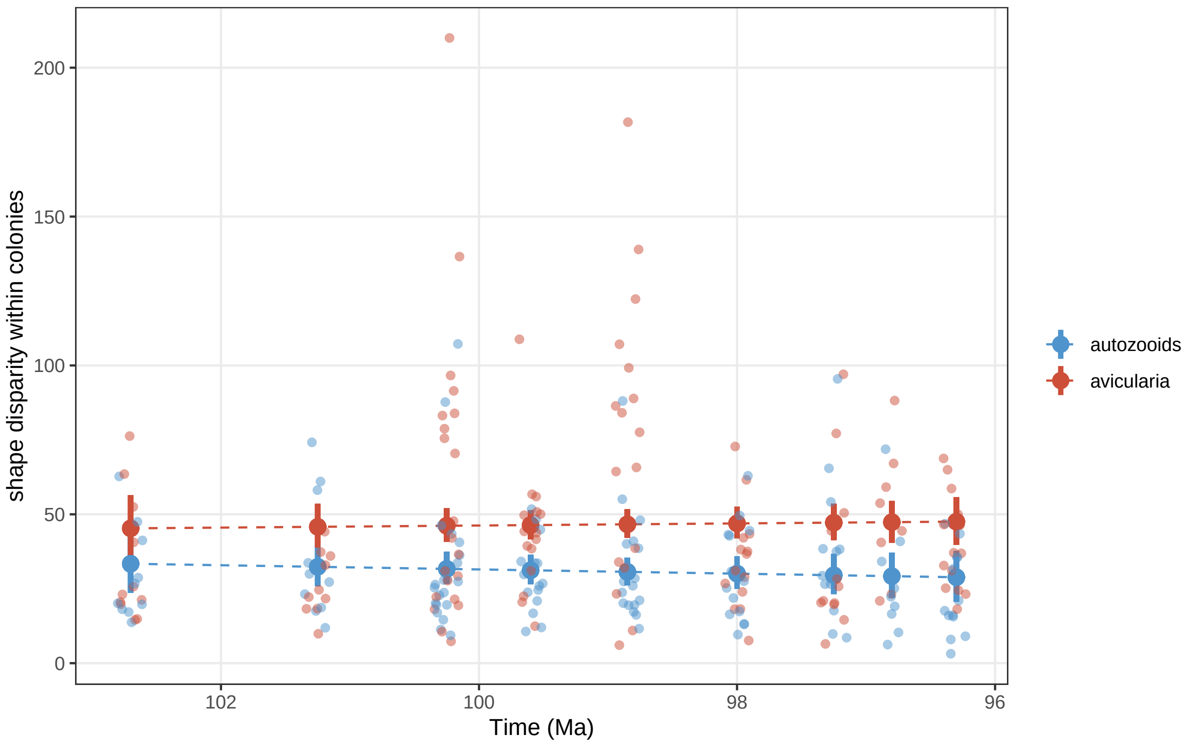

Figure S8: Disparity within colonies for each zooid type and across time intervals. Disparity values for colonies are shown as transparent points, with mean predicted disparity values from model 6 are shown as filled opaque points and 95% confidence intervals shown as lines around each mean point. This pattern mirrors the among-colony pattern in Fig. S7, where within colony disparity of avicularia is higher on average than autozoid disparity through time (Table S6).

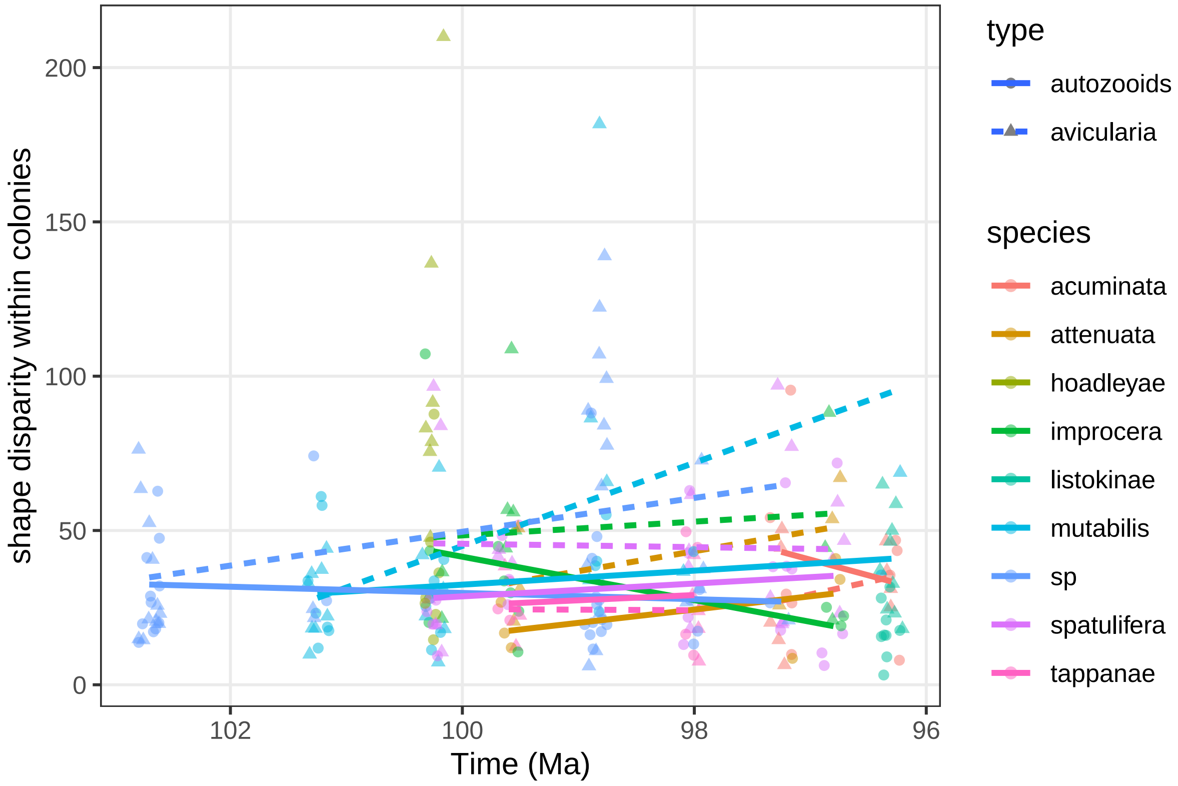

Figure S9: Disparity within colonies over each time interval and across species.

functions can evolve, which makes it difficult to infer original functionality based on extant function. Nevertheless, it is possible to indirectly infer zooid functionality from features preserved in the zooid skeleton: specifically, the appearance of ovicells on zooids. The presence of ovicells on the distal end of a zooid indicates that the zooid possesses a functioning feeding apparatus (lophophore) as the tentacles of the lophophore contain the intertentacular organ that produces gametes (17, 18, 19).

In most extant cheilostomes, autozooids have fully developed lophophores and can produce ovicells, while avicularia have heavily vestigialized polypides that cannot support lophophores or produce ovicells (15, 20, 21). Thus, ovicell-budding capabilities reflect the functional differences between autozooids and avicularia that occur as a result of modification to soft-tissue morphology (15, 20, 22). Furthermore, in the few extant taxa with subtly morphologically differentiated avicularia, like some species of *Callopora*, avicularia retain their ovicell-budding capabilities (23), suggesting that functional differentiation does not occur until avicularia become sufficiently divergent from autozooids.

Like in *Callopora*, some *Wilbertopora* avicularia retain the ability to produce ovicells (5, 23, 24), indicating that early avicularia were likely functionally continuous with autozooids. While the precise functional capabilities of these avicularian ovicells has been questioned, their appearance in certain defined species of *Wilbertopora* is thought to be functional due to the visibility of the ovicell opening. We follow Cheetham et al. 2006 in the determination of which avicularian ovicells are functional by species identification. The loss of ovicell-budding capabilities can therefore be an indication of functional differentiation in avicularia, since reduction of the polypide will necessarily impede reproductive function and feeding ability (9, 24).

High morphological disparity of avicularia may also be associated with their loss of autozooidal functions. High ranges of morphological variation is a common feature of vestigiality (25, 26), which has been used to explain the high levels of variation observed in some extant cheilostome with highly derived, seemingly multipurpose avicularia (21, 27). In *Wilbertopora*, high morphological disparity of avicularia relative to autozooids in colonies may indicate that polypides have become vestigial and are no longer functional in feeding or reproduction.

High degrees of avicularian divergence from autozooid morphology in colonies may also reflect functional differentiation of avicularia, since autozooid morphology is known to be constrained by geometric constraints of supporting a fully functioning polypide (15). Avicularia that are highly divergent from autozooids have presumably overcome this morphological constraint, indicating potential loss of polypide functionality.

Due to the vestigial nature of lophophores in functionally divergent avicularia in extant taxa, it is expected that functionally distinct avicularia in *Wilbertopora* will have more disparity and divergence from autozooids within colonies than functionally continuous avicularia. While the absence of avicularian ovicells in colonies does not necessarily indicate that avicularia lack the ability to generate ovicells, we can view the presence of avicularian ovicells as an indication of functionally indistinguishable avicularia and autozooids. Using the presence of avicularian ovicells as our null condition/state, we test whether colonies lacking avicularian ovciells have more divergence and more disparity than those possessing avicularian ovicells. If colonies with high divergence and high disparity tend to lack avicularian ovicells, then such colonies may possess avicularia that have vestigialized polypides. Furthermore, if ovicell frequency is significantly reduced in these colonies, then there is evidence for colony-level specialization, which would suggest that different zooid types serve different functions in colonies.

#### 7.2 Collection of ovicell data

Colonies with ovicells were checked for the presence of ovicells on avicularia using meshes and photographs. Colonies that lack ovicells or fall below a certain size (<30 zooids) were excluded from this portion of the analysis because smaller colonies may not have reached sexual maturity yet. For each colony, all zooids were counted to estimate colony size, and ovicells were counted to estimate ovicell frequency.

#### 7.3 Test statistics for significant differences.

Linear modeling was used to determine whether there is a significant relationship between divergence within colonies and the presence or absence of avicularian ovicells in colonies (model 7, Table S7).

**Model 7:** $lm.rrpp(DST \sim ovicells)$

where *DST* is the distance between zooid types in colonies, and *ovicells* is the presence or absence of avicularian ovicells in colonies.

Table S7: ANOVA for regression of distances between mean orifice and opesia shapes of zooid types within colonies as a function of avicularian ovicell status. Table header as in Table S1.

| Terms | Df | SS | MS | $R^2$ | F | Z | Pr>F |
|---|---|---|---|---|---|---|---|
| *Ovicells* | 1 | 310 | 310 | 0.21 | 18 | 3.1 | 0.001 |
| Residuals | 66 | 1100 | 17 | 0.79 | | | |
| Total | 67 | 1400 | | | | | |

Colonies lacking avicularian ovicells have significantly more divergent zooid types in colonies than those possessing avicularian ovicells.

Linear modeling was also used to determine whether there is a significant relationship between avicularian disparity within colonies and the presence of avicularian ovicells within colonies (model 8, Table S8).

**Model 8:** $lm.rrpp(Disp_{avic} \sim ovicells)$

Where $Disp_{avic}$ is the disparity of avicularia within colonies, and *ovicells* is the presence or absence of avicularian ovicells in colonies.

Table S8: ANOVA for regression of avicularian disparity within colonies as a function of avicularian ovicell status. Table header as in Table S1.

| **Terms** | **Df** | **SS** | **MS** | **$R^2$** | **F** | **Z** | **Pr>F** |
|---|---|---|---|---|---|---|---|
| *Ovicells* | 1 | 1700 | 1700 | 0.038 | 2.6 | 1.2 | 0.12 |
| Residuals | 66 | 44000 | 660 | 0.96 | | | |
| Total | 67 | 45000 | | | | | |

The model results suggest that mean values of avicularian disparity is on the margin of significantly different between colonies that possess and lack avicularian ovicells.

To estimate whether ovicell frequency differs between colonies that possess ovicells on avicularia and those that lack them, ovicell frequency was regressed against the status of avicularian ovicells (present, absent) in colonies:

**Model 9:** $lm.rrpp(ovicellfrequency \sim ovicells)$

Table S9: ANOVA for regression of ovicell frequency within colonies as a function of avicularian ovicell status. Table header as in Table S1.

| **Terms** | **Df** | **SS** | **MS** | **$R^2$** | **F** | **Z** | **Pr>F** |
|---|---|---|---|---|---|---|---|
| *Ovicells* | 1 | 0.12 | 0.12 | 0.076 | 4.6 | 1.7 | 0.039 |
| Residuals | 56 | 1.5 | 0.027 | 0.92 | | | |
| Total | 57 | 1.6 | | | | | |

There is a significant difference in ovicell frequency between groups.

### 8 Modeling diffusion of energy within a colony

Let's assume $n$ connected nodes (neighboring physiologically connected zooids), with $m$ edges. An edge exists between two nodes if they share a border.

Let $A$ be the $n \times m$ incidence matrix, such that

$$A_{ij} = \begin{cases} -1, & \text{edge } j \text{ points from node } i \\ +1, & \text{edge } j \text{ points to node } i \\ 0, & \text{otherwise.} \end{cases}$$

Also, let $f$ be the flow vector (e.g., of nutrients from zooid to zooid along the funicular system) of length $m$, and let $s$ be the source vector of length $n$.

***Flow conservation*** across the graph is represented by the following $n$ equations:

$$Af + s = 0. \tag{1}$$

Let $v$ be the potential vector of length $n$, and $R$ be the $m \times m$ resistance matrix. We'll assume this is a diagonal matrix $R = \text{diag}(r)$, for some vector $r$, where $r_i$ is the resistance along edge $i$.

In a ***diffusion system***, the flow along an edge is proportional to the potential difference across adjacent nodes ($m$ equations):

$$Rf = -A^T v. \tag{2}$$

Now, let's fix the potential of each node. Let $\mathcal{P}$ be the set of indices of feeders, and $\mathcal{P}^c$ (i.e., the complement of $\mathcal{P}$) be the set of indices of non-feeders. We assume

$$v_i = 1, i \in \mathcal{P}$$

$$v_i = 0, i \in \mathcal{P}^c$$

We ***specify the potential*** with a (trivial) matrix-vector equation (n equations):

$$Iv = w \tag{3}$$

where $I$ is the identity matrix, and $w$ collects the values above (+1 or 0 for feeders and non-feeders, respectively). (This equation leaves $v$ as a variable to solve for, but sets the potentials equal to whatever one wants in $w$.)

These three equations can be combined into a single statement using block matrices:

$$\begin{bmatrix} A & I & 0 \\ R & 0 & A^T \\ 0 & 0 & I \end{bmatrix} \begin{bmatrix} f \\ s \\ v \end{bmatrix} = \begin{bmatrix} 0 \\ 0 \\ w \end{bmatrix}, \tag{4}$$

which we can also write as

$$Kx = y.$$

Then the solution we are looking for is $x = K^{-1}y$. This solves for the flow vector $f$ and the source vector $s$ (and keeps $v = w$), according to the constraints above.

We interpret the source vector $s$ biologically as describing the pattern of the dearth or excess of energy for each zooid within the colony. Negative elements of $s$ represent the energy sinks of non-feeding members. Elements that are either positive or 0 represent feeding members of the colony. And the positive elements of $s$ we interpret as the excess energy required to support non-feeding members. Conditioned on the fact that the colonies we quantify were alive with the network we estimate $s$ with, we can estimate three quantities: (A) the excess energy acquired by the colony and (B) the average energetic benefit per zooid. And (C) the average energetic cost of non-feeding members. We define A to be the sum of all non-negative elements of $s$. And B to be the average of all non-negative elements of $s$. And C to be the average of all negative elements of $s$.

The specific values of the potential vector $v$ and the resistance matrix $R$ can be modified with this approach. We fixed them here to make all colonies comparable. For resistance we set $r_i = 1$ for all zooids which represents a biological assumption that all zooids have equal ability to move or receive energy from neighbors. For the potentials, we set all feeders to equal 1 and all non-feeders to equal 0. One can envision alternative schemes, where perhaps vicarious avicularia have larger potentials than do adventitious avicularia because vicarious avicularia can bud asexually while the latter cannot.
